# Interparental Gene Conversion in General Population: A Novel Mechanism For Loss of Heterozygosity

**DOI:** 10.1101/2024.12.05.626979

**Authors:** Jumpei Toratani, Masahito Tachibana, Junichi Sugawara, Atsushi Sugawara, Takeki Sato, Yuri Takahashi, Hiroaki Hiraga, Emi Yokoyama, Zen Watanabe, Masatoshi Saito, Nobuo Yaegashi, The Tohoku Medical Megabank Project Study Group, Gen Tamiya, Jun Takayama

## Abstract

Gene conversion is a process in which genetic material from a donor sequence is unidirectionally copied to an acceptor sequence during the homologous recombination repair of a DNA double-strand break. Although gene conversion has been widely studied in the context of meiosis, hereditary diseases, and cancer development, gene conversion between parental homologs in the zygotes remains controversial. Here, we developed a method to detect interparental gene conversions by focusing on Mendelian errors and identified gene conversion events in one out of every 21.8 births. Some of these events were observed in genetic regions, potentially affecting offspring phenotypes. Interparental gene conversion leads to the offspring inheriting two identical alleles from one parent, resulting in a loss of heterozygosity. Our findings suggest that naturally occurring interparental gene conversions may provide a novel mechanism for the development of certain genetic diseases.

## Introduction

Gene conversion is a process in which a portion of the ‘donor’ DNA sequence is copied unidirectionally to the ‘acceptor’ DNA sequence during the repair of a DNA double-strand break (DSB) (Chen et al. 2007). DSBs are commonly repaired via non-homologous end-joining (NHEJ) or homologous recombination (HR) (Vítor et al. 2020). NHEJ joins two DNA ends with minimal reference to the DNA sequence (Shrivastav et al. 2008; Scully et al. 2019). In contrast, HR employs homologous regions as a template, and as a result, it can lead to gene conversion (Chen et al. 2007; Sun et al. 2020). In meiosis, gene conversion can lead to non-Mendelian inheritance, which drives genome evolution by creating new allelic combinations; however, this process is also known to cause genetic diseases. On the other hand, in mitosis, if HR employs homologs as templates, the converted regions can become homozygous. Consequently, gene conversion causes a loss of heterozygosity (LOH), potentially leading to cancer development (Chen et al. 2010). Gene conversion may also occur shortly after fertilization as a repair process for DSBs formed in gametes or zygotes (Wilde et al. 2021; Liang et al. 2023). In such cases, the DNA sequence from one parent serves as the repair template, resulting in an offspring with two copies from that parent. However, some researchers question the existence of such gene conversion (Egli et al. 2018), given that the parental genomes remain physically separated into two pronuclei until just before the first cell division stage.

Several factors can cause DSBs. In meiosis, most DSBs are observed at hotspots determined by certain proteins (Kong et al. 2010; Pratto et al. 2014; Halldorsson et al. 2019). In mitosis, DSBs are triggered by radiation, stalled replication forks, or specialized endonucleases. Recently, the advent of CRISPR/Cas9 has enabled the targeted generation of DSBs. Using this technique, several studies have attempted to edit the genes of preimplantation embryos via homology-directed repair (HDR) (Wu et al. 2013; Zhang et al. 2023). In a study by Ma et al. (Ma et al. 2017), the CRISPR/Cas9 was designed to target paternal mutant alleles, and the Cas9 protein was injected during fertilization. These DSBs were repaired via HDR using maternal homologs as templates, resulting in gene conversion that we hereafter refer to as interparental gene conversion. Additionally, gene conversion during embryogenesis is longer than in somatic cells, with a tract length of approximately 20 kb (Donoho et al. 1998; Mansai et al. 2011; Liang et al. 2023). Extensive interparental gene conversion can lead to genetic disorders due to the homozygosity of deleterious alleles or lead to the development of imprinting abnormalities by erasing epigenetic DNA modifications. Hence, utilizing gene conversion as a tool for gene therapy remains impractical at this time (Liang et al. 2023). Nevertheless, these findings indicate that in cases where DSBs spontaneously arise in gametes or zygotes, interparental gene conversion can occur, resulting in the offspring inheriting two copies from one parent. This might explain the mechanisms underlying a subset of recessive genetic diseases and imprinting disorders whose origins remain obscured until now. Although gene conversion has been widely studied in the context of meiosis, hereditary diseases, and cancer development, the frequency and genetic impact on offspring of naturally occurring interparental gene conversion have yet to be evaluated.

In this study, we aimed to develop a pipeline employing whole-genome sequencing data from 2,302 Japanese parent-offspring trios recruited from the TMM BirThree Cohort Project (Kuriyama et al. 2020) to examine the existence of naturally occurring interparental gene conversion. All trios were analyzed with short-read whole-genome sequencing data, and among them, 109 trios were additionally analyzed using nanopore long-read whole-genome sequencing data. Our results suggest that spontaneous DSBs in gametes or zygotes are repaired using the DNA sequence of the other parent as a template around the time of fertilization, leading the offspring to inherit two copies from one parent. These findings provide new insights into genetic disorders and contribute to advancements in reproductive medicine.

## Results

### Cohort Characteristics

Our study targeted 2,302 trio families confirmed to have biological parent-child relationships, each with whole-genome sequencing data obtained through short-read technology. The ages of the participants ranged from 0 to 82 years, with median ages of 62, 32, and 0 years across three generations, respectively (Supplemental Fig. S1). A total of 45,482,713 single nucleotide variants (SNVs) were identified in all samples. After applying various quality control (QC) filters (Methods), the average number of targeted SNVs was 3,248,591 ± 277,792 per trio family.

### Pipeline for Identifying Interparental Gene Conversion

We developed a pipeline to identify interparental gene conversion by focusing on LOH, especially non-Mendelian LOH, in the offspring genotype (Fig. 1). In our autosomal whole-genome sequencing data, we targeted biallelic single nucleotide variants (SNVs). When focusing on a single position, considering that each individual can be in one of three states— reference homozygous, alternative homozygous, or heterozygous—27 possible genotype combinations for the trios exist. We classified genotype combinations into three categories: sites consistent with gene conversion with Mendelian errors, sites inconsistent with gene conversion, and sites consistent with gene conversion but without Mendelian errors (Supplemental Table S1). The first and second categories were extracted as gene conversion sites and non-gene conversion sites, respectively (Fig. 1B). The third category, which includes cases in which all trio members are homozygous reference, was not employed as a marker because it does not specifically indicate gene conversion. Following previous studies (Chen et al. 2007; Liang et al. 2023), we defined minimum gene-conversion regions as regions enclosed by at least two consecutive gene conversion sites without any non-gene conversion sites and maximum gene-conversion regions extending to the nearest non-gene conversion sites beyond the minimum gene-conversion regions (Fig. 1A). Each interparental gene conversion region was required to contain at least two consecutive Mendelian errors consistent with gene conversion. While a single Mendelian error can occur due to mapping or variant calling errors, requiring consecutive Mendelian error sites is expected to significantly decrease the occurrence of false positives.

**Figure 1.**
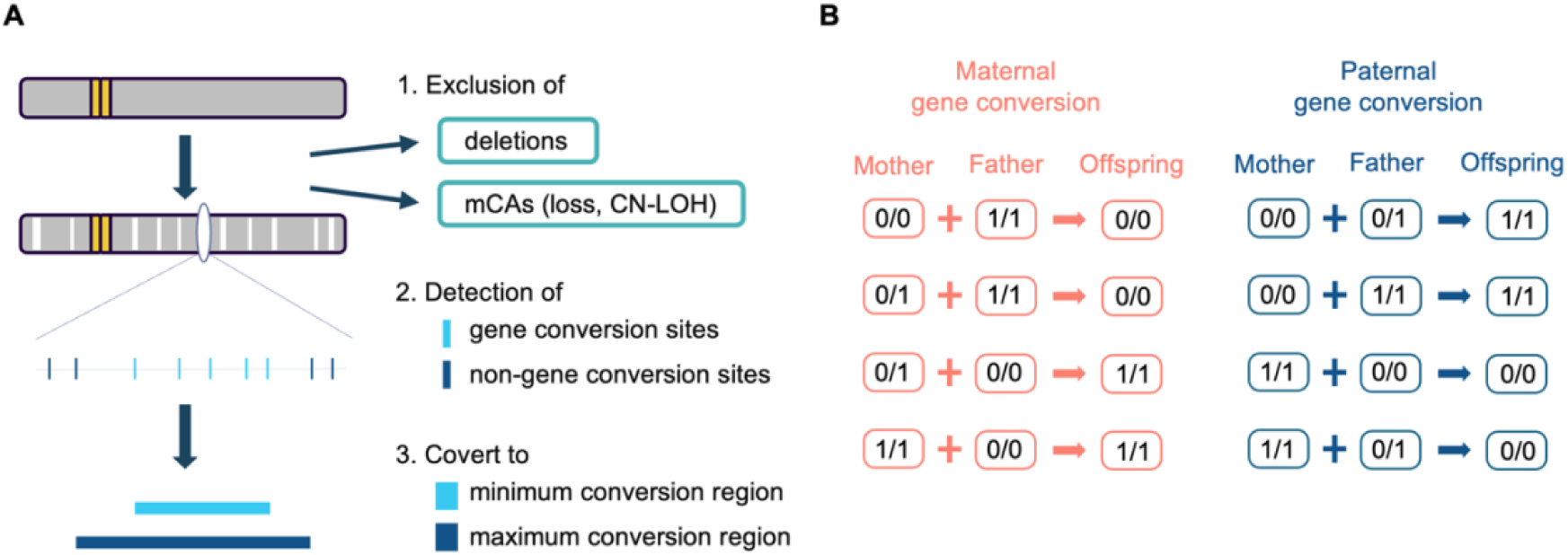
Overview of the detection method for interparental gene conversion. *(A)* Pipeline of identifying interparental gene conversion by focusing on non-Mendelian LOH shown in the offspring’s genotype. *(B)* Genotype combinations extracted as gene conversion sites. These combinations are consistent with gene conversion and exhibit Mendelian errors. 0/0 means homozygous reference, 1/1/ means homozygous alternative, and 0/1 means heterozygous state.

This formed the core of our method for identifying interparental gene conversions. However, several factors must be considered: (1) deletion, (2) uniparental disomy (UPD), and (3) mosaic chromosomal alteration (mCA). These considerations are necessary because deletions and UPD lead to LOH (Landsverk et al. 2012), making it essential to differentiate them from interparental gene conversion. Additionally, mCAs can cause the DNA sample to not accurately reflect the germline (Terao et al. 2020).

Although deletion, UPD, and interparental gene conversion all result in an offspring inheriting only one parent’s allele, there are subtle differences in the genotypes of the trio (Supplemental Fig. S2). Deletion results in only one copy of the affected region, whereas both UPD and interparental gene conversion result in two copies. UPD typically originates from non-disjunction in gametogenesis or postzygotic recombination, spanning an entire chromosome or a large segment of it (Kotzot 2001; Niida et al. 2018). Hence, studies aiming to identify UPD often set certain lengths as criteria (Nakka et al. 2019; Scuffins et al. 2021). Based on previous studies, we set two criteria for defining UPD: the inclusion of telomeres and a minimum length of 5 Mbp.

mCA is an aging-related somatic mutation detectable in peripheral blood, characterized by the clonal expansion of structural changes (Terao et al. 2020; Jakubek et al. 2023a; Jakubek et al. 2023b). Because specimens affected by mCAs do not represent the germline genotype, this phenomenon can distort the results of studies which focus on Mendelian errors. As part of our quality control, we calculated the heterozygosity for each sample (heterozygous sites/total sites) and identified samples with significantly decreased the heterozygosity, particularly in elderly individuals (Supplemental Fig. S3). Furthermore, we confirmed a negative correlation between the length of mCAs and the heterozygosity (Supplemental Fig. S4). Based on these findings, we attributed the decrease in the heterozygosity partially to age-related increases in mCAs.

Accordingly, we identified the mCA regions that could affect genotyping and excluded them from the analysis. In particular, mCAs showing loss or copy-neutral LOH (CN-LOH) may cause heterozygous sites to be incorrectly called homozygous sites. We applied stringent QC to the depth of coverage for homozygous sites; thus, theoretically, genotyping should not be affected unless more than 90% of the cells exhibit mCAs. To minimize false positives, we adopted a stringent approach by excluding regions from this analysis where over 50% of the cells showed mCAs involving loss or CN-LOH.

Considering these points, we developed a pipeline to identify interparental gene conversions using short-read whole-genome sequencing data (Fig. 1A). First, we identified deletions and mCAs showing loss or CN-LOH in every sample of the trio and excluded them from the analysis. For the remaining regions, we extracted gene conversion sites and non-gene conversion sites. Finally, using these sites, we identified minimum gene-conversion regions and maximum gene-conversion regions.

### Detection of Uniparental Disomy (UPD)

This study also enables the detection of UPDs, as both interparental gene conversion and UPD result in offspring inheriting two copies from one parent. The UPDs can be categorized into four types (Supplemental Fig. S5) (Scuffins et al. 2021). We identified two whole-chromosome UPDs (isodisomy and mixed UPD) and one segmental UPD (Fig. 2). No individuals had UPDs affecting more than one chromosome. A trio named UPD-1 exhibited maternal heterodisomy on chromosome 22. A trio UPD-2 exhibited a maternal mixed UPD on chromosome 8, which was divided into three segments: two isodisomies and one heterodisomy. This mixed UPD was hypothesized to result from two recombination events on the maternal chromosome during meiosis. A trio UPD-3 exhibited a maternal segmental UPD covering most of the long arm of chromosome 2. Collectively, 3 out of 2302 trios had UPDs, aligning with previously published data (Nakka et al. 2019; Scuffins et al. 2021). Both whole-chromosome UPDs originated from the mother, which is consistent with previous findings that whole-chromosome UPDs are mainly attributed to chromosomal non-disjunction and are therefore predominantly maternal (Scuffins et al. 2021). Three trios identified with UPDs were excluded from subsequent analysis of interparental gene conversion.

**Figure 2.**
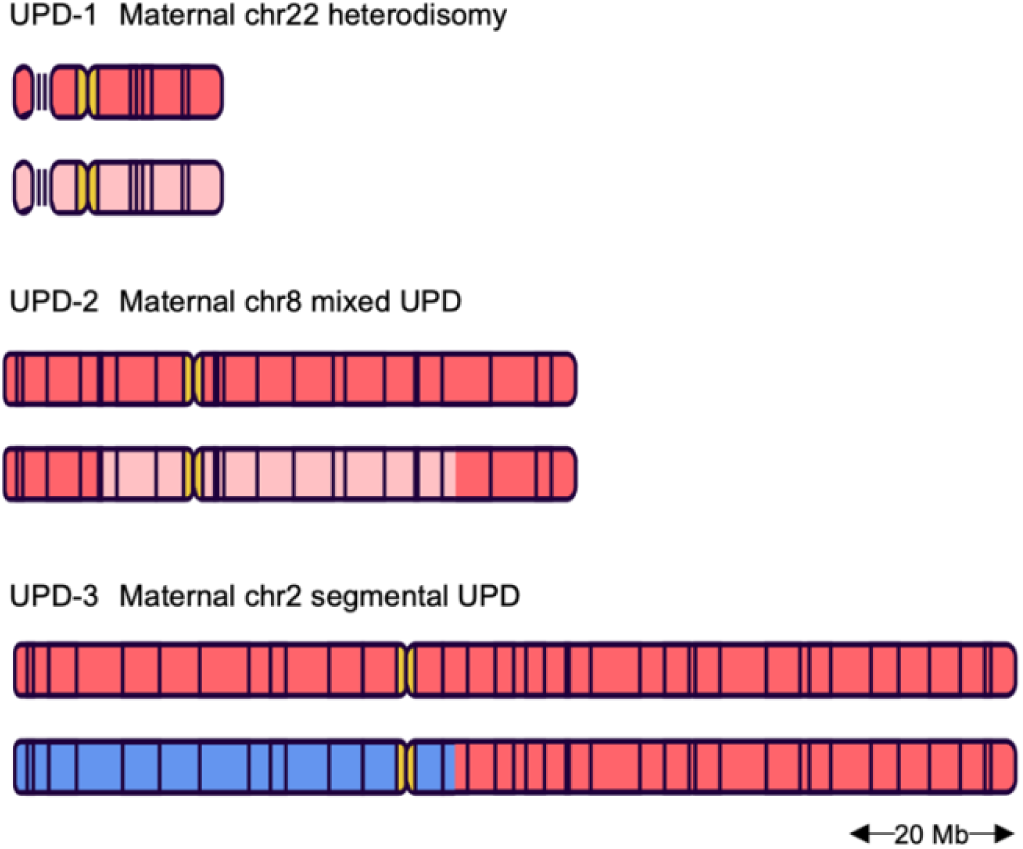
Schematic illustration of 3 UPDs identified in 2302 trios. UPD-1 exhibited maternal heterodisomy on chromosome 22. UPD-2 exhibited maternal mixed UPD observed on chromosome 8, divided into three segments: two isodisomies and one heterodisomy. UPD-3 has a segmental UPD covering most part of the long arm of chromosome 2. The dark red and light red each represent maternal chromosomes, while the blue represents paternal chromosomes. This illustration is adapted from TogoTV (© 2024 DBCLS TogoTV, CC-BY-4.0).

### QC using long-read technology

We verified the interparental gene conversion using long-read sequencing. Using only short-read technology, we identified 258 maternal and 248 paternal gene conversions among 2299 trios (Supplemental Fig. S6-S8). The frequency of interparental gene conversion per sample was 0.112 for maternal origins and 0.108 for paternal origins, with no significant difference observed between maternal and paternal origins (Z-test for proportions). Our workflow for detecting interparental gene conversions involved extracting deletions to ensure that they were not misidentified as gene conversions. However, short-read technology has limitations in sensitively detecting structural variants (SVs), including deletions, due to the difficulty of spanning repetitive sequence (Kosugi et al. 2019; Belyeu et al. 2021; Lei et al. 2022). In fact, recent comparisons between short-read and long-read technologies have demonstrated that SVs are frequently overlooked with short-read technology (Chaisson et al. 2019; Zhao et al. 2021). Owing to limitations of short-read technology, overlooked deletions may be mistakenly identified as interparental gene conversions. Therefore, we evaluated this possibility using SV calls made with long-read technology in a subset of 109 out of the 2299 trios. We initially identified 12 maternal and 9 paternal gene conversions in these 109 trios using only short-read technology. Employing long-read technology, we discovered that five of the maternal gene conversions (41.7%) and two of the paternal gene conversions (22.2%) were deletions within the children’s sequences (Fig. 3).

**Figure 3.**
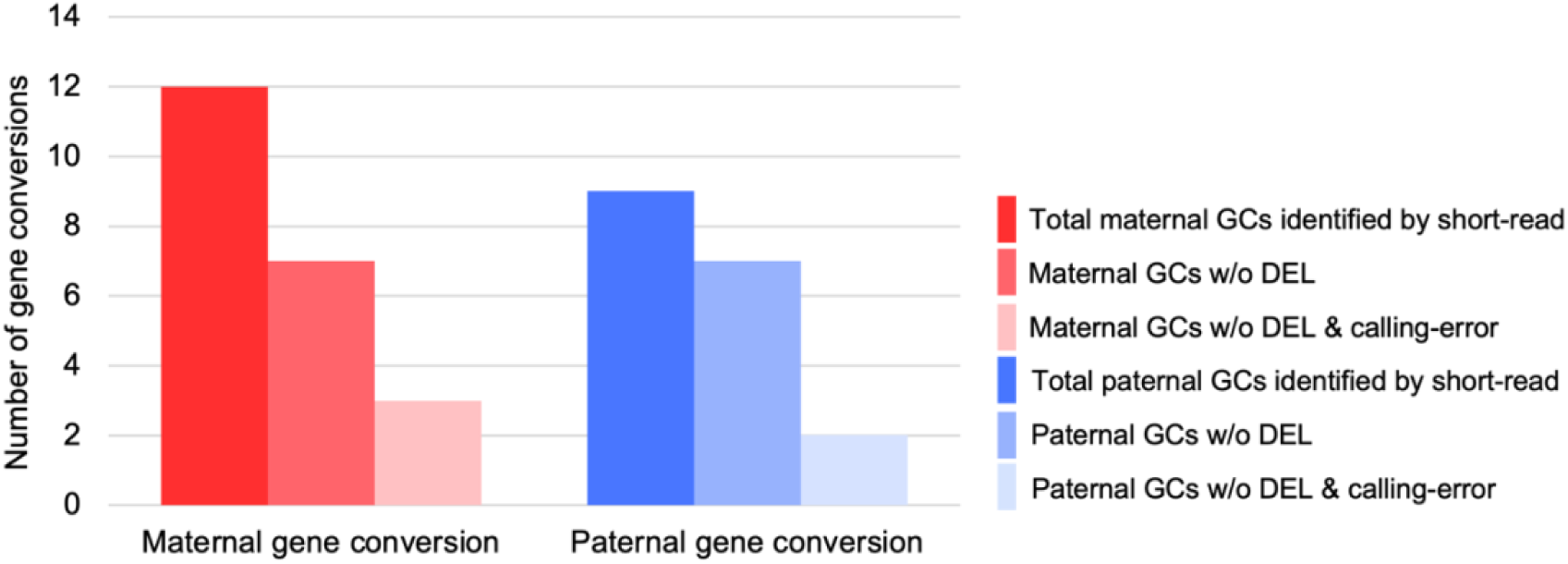
Validation of interparental gene conversion using long-read technology. Using short-read technology, 12 maternal and 9 paternal gene conversions were identified in 109 trios. 5 of the maternal gene conversions and 2 of the paternal gene conversions were actually deletions within the children’s sequences. 4 maternal cases and 5 paternal cases were judged to be calling errors caused by short-read technology. As a result, 3 out of 12 maternal and 2 out of 9 maternal gene conversions were consistently validated by long-read technology. ‘GC’ stands for gene conversion.

Since we identified interparental gene conversions by relying on Mendelian errors, errors in SNV calling could lead to false positives. Stringent quality controls were used to minimize false positives. Despite these measures, errors at various stages such as DNA extraction, sequencing, and mapping can lead to inaccurate SNV calling. To assess this possibility, we employed long-read sequencing data to verify the remaining 14 gene conversions, which had been confirmed not to be deletions. Each gene conversion was visually inspected by two specialists using the Integrative Genomics Viewer (IGV) (Robinson et al. 2011). Consequently, four maternal and five paternal cases were not confirmed as gene conversions, suggesting these were likely errors originating from short-read technology. Nearly 80% of interparental gene conversions identified by short-read technology were either deletions or calling errors (maternal: 9 out of 12, 75%; paternal: 7 out of 9, or 77.8%) (Fig. 3). Nevertheless, over 20% of cases were consistently validated using long-read technology, providing strong evidence for naturally occurring interparental gene conversions. In summary, using long-read technology in combination with short-read technology, we verified the presence of three maternal and two paternal gene conversions in 109 trios. Taken together, we conclude that interparental gene conversions occurred at a frequency of one in every 21.8 births, with maternal gene conversions at one in every 36.3 births and paternal gene conversions at one in every 54.5 births.

Next, we annotated these events to genomic features to evaluate the potential effects of interparental gene conversions observed in the five trios (Fig. 4). In trio named GC-1, maternal gene conversion was observed on chromosome 5p13.1, with a minimum length of 9,372 bp and a maximum length of 59,215 bp. This region includes *ZDHHC11B*, which encodes a protein involved in palmitoylation, a process crucial for membrane association and protein function (Dai et al. 2023). In trio GC-2, maternal gene conversion was identified on chromosome 14q12, with a minimum length of 67,662 bp and a maximum length of 160,980 bp. The gene conversion region overlapped with *MIR3171HG*, which has been implicated in various RNA-related processes (Pruitt et al. 2014). In trio GC-3, maternal gene conversion was observed on chromosome 22q11.23-q12.1, with a minimum length of 244,086 bp and a maximum length of 276,284 bp. Two pseudogenes, *IGLL3P* and *CRYBB2P1*, are located within the gene conversion region. In trio GC-4, paternal gene conversion was identified on chromosome 4q13.2-q13.3, with a minimum length of 2,911 bp and a maximum length of 146,266 bp. The maximum gene conversion region covered *UGT2B28*, which is involved in steroid hormone metabolism (Lacombe et al. 2023). In trio GC-5, paternal gene conversion was identified on chromosome 10q11.21, with a minimum length of 42,267 bp and a maximum length of 450,123 bp. None of the genes were affected by this event. Collectively, among the five gene conversions, two altered the genotype within the gene regions, indicating potential effects on the phenotypes of the offspring. None of the detected interparental gene conversions were located in regions associated with imprinting disorders (Butler 2020; Beygo et al. 2023).

**Figure 4.**
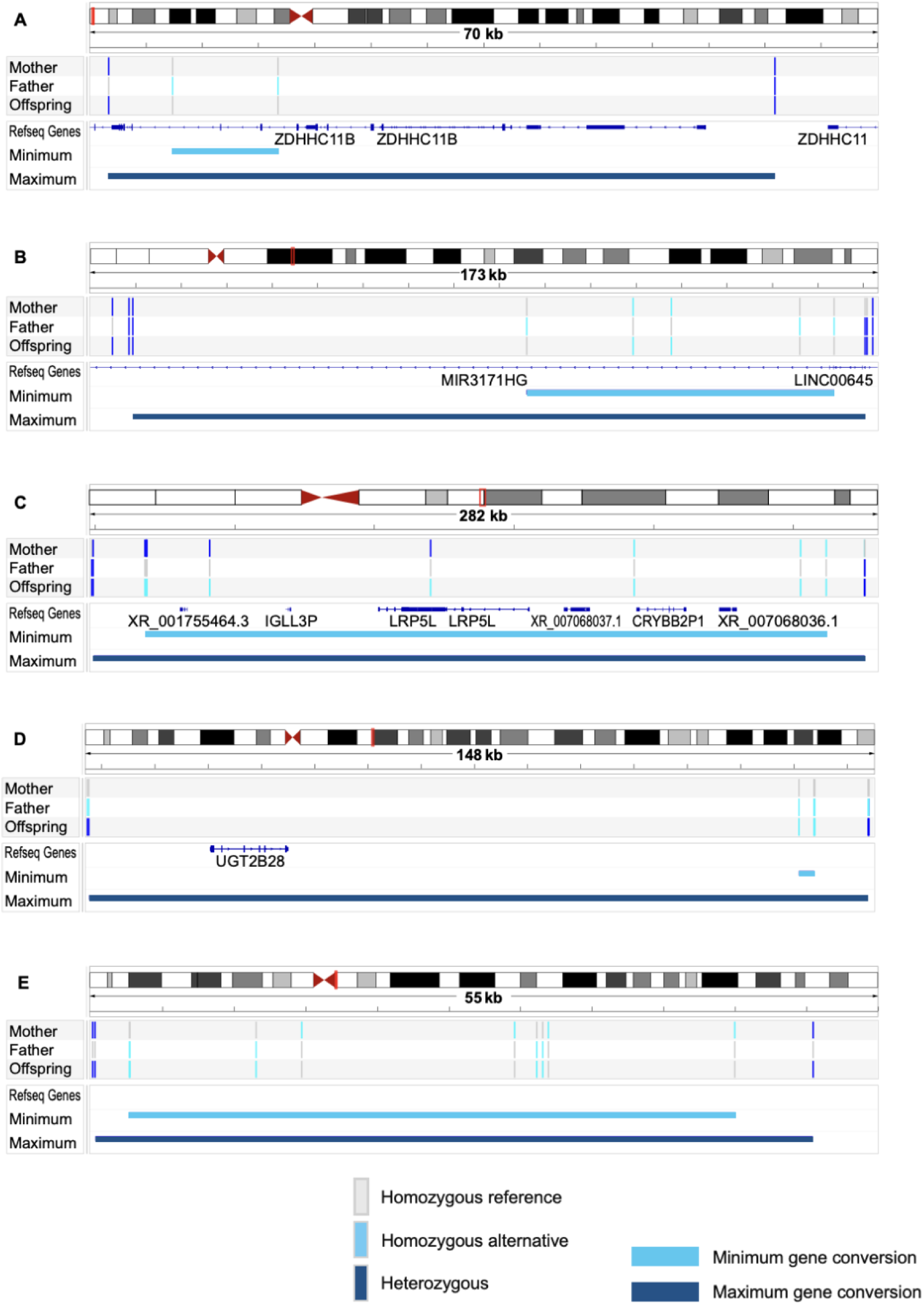
Five interparental gene conversions validated via long-read technology. Each small box shows the genotype of a family member, with the mother at the top, the father in the middle, and the child at the bottom. (A) Maternal gene conversion on chromosome 5p13.1 (GC-1). (B) Maternal gene conversion on chromosome 14q12p (GC-2). (C) Maternal gene conversion on chromosome 22q11.23-q12.1 (GC-3). (0) Paternal gene conversion on chromosome 4q13.2-q13.3 (GC-4). (£) Paternal gene conversion on chromosome 10q11.21 (GC-5)

The origins of these interparental gene conversions are DSBs naturally occurring around fertilization. One plausible situation where DSBs may arise in gametes is during meiosis in gametogenesis. DSBs occurring during meiosis might persist until fertilization and be repaired through homologous recombination using the other parent’s allele as a template, resulting in interparental gene conversion. To evaluate this possibility, we assessed the relationship between DSB hotspots during meiosis (Takayama et al. 2024) and the interparental gene conversions observed in our study. Among the five interparental gene conversions, one had a recombination hotspot within the maximum gene conversion region. Hence, while DSBs occurring during meiosis may contribute to interparental gene conversion, other factors may also be involved.

## Discussion

In this study, we developed a method to detect interparental gene conversions by focusing on Mendelian errors and successfully identified gene conversions at a rate of one in every 21.8 births. Although several studies using CRISPR/Cas9 to intentionally create DSBs have confirmed gene conversion between parental alleles during early embryonic development, to the best of our knowledge, this is the first study to identify interparental gene conversion in the general population and evaluate its frequency.

Interparental gene conversion is a result of interhomolog repair. While interhomolog repair is commonly observed during meiosis, it is rarely seen in somatic cell division, and whether it occurs in zygotes is controversial due to the physical separation of parental genomes into separate pronuclei (Wilde et al. 2021). Nevertheless, a study that used time-lapse light-sheet microscopy to track mitosis in mouse zygotes revealed a window of approximately 45 min after nuclear envelope breakdown, during which the maternal and paternal genomes overlapped (Reichmann et al. 2018). Indeed, in zygotes during the late S/G2 phase, endogenous DSBs can originate, and they can be repaired via interhomolog gene conversion (Wilde et al. 2021). Given that most of the interparental gene conversions identified in our study were not located in meiotic recombination hotspots, these DSBs may have originated naturally at the time of fertilization.

Although the frequency of gene conversion during meiosis is well understood, its occurrence in embryos remains unclear. In a study using mice, *RAD51* puncta were observed in approximately 20% of the fertilized eggs in the late S/G2 phase (Wilde et al. 2021), suggesting that interparental gene conversion could potentially occur at this frequency. We identified interparental gene conversion at a frequency of about 5%. To reduce false positives, we required each gene conversion to have at least two consecutive Mendelian errors. Consequently, smaller events, or events that did not exhibit Mendelian errors, could not be identified. Therefore, the calculated frequency is likely underestimated. Considering that male gametes are more vulnerable to damage and prone to DSBs than female gametes (Derijck et al. 2008; Ménézo et al. 2010), it can be inferred that maternal gene conversions are more frequent than paternal ones. Additionally, intracytoplasmic sperm injection (ICSI), a technique of assisted reproductive technology (ART), may induce the transgenesis of foreign genes (Moreira et al. 2005), potentially resulting in an increase in interparental gene conversion in offspring born via this technique. The subset analyzed using long-read technology did not include any individuals born via ART, and the results obtained using only short-read technology did not reveal any significant differences between the methods of conception (Supplemental Fig. S9). Although we did not observe any significant impact of the sex or conception method on the frequency of interparental gene conversion, future studies with larger sample sizes could provide more accurate frequency estimates and a clearer evaluation of the impact of these factors.

Gene conversion tract length is known to differ between meiosis and mitosis. In meiosis, the tract length is less than 1 kb (Sun et al. 2012; Li et al. 2019), whereas in somatic cell division, it spans several kilobases (Neuwirth et al. 2007). The tract length of interhomolog gene conversion in zygotes or early embryos is even longer than that in somatic cells, potentially extending beyond 10 kb (Wilde et al. 2021; Liang et al. 2023). This implies that interhomolog repair in early embryos may have different characteristics compared to that in germ cells and somatic cells. We identified gene conversions with minimum tract lengths ranging from 2,910 bp to 244 kb and maximum tract lengths ranging from 50 kb to 276 kb, which were considerably longer than the gene conversion tracts induced by CRISPR/Cas9. Given this length, there could be concerns that these regions might be interstitial segmental UPD (Kotzot 2008). However, interstitial segmental UPD is remarkably less frequent than telomeric segmental UPD (Scuffins et al. 2021). Hence, it is unlikely that most of the cases identified here are interstitial segmental UPD.

UPD is defined as the inheritance of both copies of a chromosome from one parent, either in its entirety or in segments (Engel 1980). Whole-chromosome UPD is primarily attributed to chromosomal non-disjunction, whereas segmental UPD arises mainly from the recombination between parental chromosomes after fertilization (Ledbetter and Engel 1995; Niida et al. 2018). Since the 2000s, several studies have investigated the prevalence of UPD in large-scale cohorts (Nakka et al. 2019; Yauy et al. 2020; Scuffins et al. 2021). A study using SNP data from the general population, focusing on identical-by-descent to identify UPD, found a prevalence of approximately 1 in 2,000 births (Nakka et al. 2019). Another study examined the incidence of UPD in children with neurodevelopmental disorders or other symptoms using trio exome SNP data and reported an incidence rate of approximately three cases per 1,000 births (Scuffins et al. 2021). However, no studies have explored the prevalence of UPD in the general population using whole-genome sequencing. In this study, the prevalence of UPD in the general population was estimated to be approximately 1.3 in 1,000 births. This frequency was approximately 2.7 times higher than that observed in a previous study that similarly targeted the general population (Nakka et al. 2019). We believe that our method, which focuses on trio Mendelian errors and utilizes whole-genome sequencing data, improves the sensitivity of this analysis.

The short-read and long-read technologies have distinct characteristics. Short-read technology is less likely to span the breakpoints of SVs, resulting in fewer detectable SVs than long-read technology (Otsuki et al. 2022). Indeed, recent long-read WGS studies have identified over 20,000 SVs per individual (Beyter et al. 2021; Wu et al. 2021), whereas short-read WGS has detected only 4,405–7,439 SVs (Eichler 2019; Collins et al. 2020). In contrast, short-read technology has higher accuracy for SNV calling (Linde et al. 2023). Hence, we extracted gene conversion sites using SNV data obtained from short-read sequencing. To minimize false positives, we applied stringent QC filters and excluded deletions and mCAs that could potentially be misidentified as interparental gene conversions. Despite these precautions, long-read technology considered approximately 80% of the identified events to be either deletions or calling errors. This highlights the limitations of short-read technology in detecting structural variants, and the potential inaccuracies that can arise from relying solely on a single method for Mendelian error extraction. Therefore, when identifying interparental gene conversions, it is recommended to use long-read technology for detecting deletions and validate results by employing another independent approach.

Our study had several limitations. First, due to ethical restrictions, we were unable to re-sequence samples or analyze additional specimens from the same individuals, which prevented us from performing experimental validation. Second, our analysis relied on data from a single tissue type, limiting our ability to completely exclude mosaicism. Despite applying stringent quality control and excluding mCAs, tissue-specific mosaic events might not be entirely ruled out. Finally, while we hypothesized that interparental gene conversions occurred around fertilization, we lacked direct evidence to confirm the timing. Further research using gametes or embryos would be necessary to provide more definitive insights regarding when this phenomenon occurs.

Taken together, we identified events suggestive of gene conversion between parental gametes in the general population and estimated its frequency. Interparental gene conversion results in offspring with two copies from one parent in the affected region, potentially leading to autosomal recessive diseases or imprinting disorders. Indeed, some cases have been reported in which despite only one parent being a carrier, the offspring develop a recessive genetic disease without any detectable deletions or UPDs (Landsverk et al. 2012). These unexplained instances of LOH can be attributed to interparental gene conversion. Thus, naturally occurring interparental gene conversion can be considered a novel mechanism for the development of genetic diseases.

## Methods

### Sample selection

We selected 2,330 Japanese parent-child trios with short-read whole-genome sequencing data from the Tohoku Medical Megabank Project Birth and Three-Generation Cohort (TMM BirThree Cohort). Trios with withdrawn consent or unverifiable biological parent-child relationships were excluded. In all neonatal samples, DNA was extracted from the umbilical cord blood. Cross-contamination from maternal DNA can be found in 2–20% of umbilical cord blood samples; however, the percentage of maternal blood contamination is typically extremely low, estimated to be approximately 10^-4 to 10^-5 of fetal nucleated cells (Hall et al. 1995; Cairo and Wagner 1997; Petit et al. 1997; Bauer et al. 2002). To assess the various types of contamination, including cross-contamination from maternal blood, we calculated the heterozygosity of all samples. Since contamination can increase the heterozygosity of a sample, we excluded two trios containing samples with the heterozygosity exceeding 34.5% (all neonates) (Supplemental Fig. S3). Based on these criteria, 2,302 trios constituted the subjects of the analysis.

### Genotyping of SNVs and structural variants

We used the genotype data from the TMM BirThree Cohort (Tadaka et al. 2023). Briefly, genomic DNA was extracted from the peripheral blood, saliva, or cord blood samples. Variant calling procedure adhered to the GATK Best Practices (DePristo et al. 2011), and reads were aligned to the GRCh38 human reference sequence. In addition to SNV calling, we used short-read sequencing data to perform SV calling using the Smoove software (Pedersen 2020), which serves as the official wrapper for Lumpy. We utilized the MoChA v1.11 caller to extract mCAs after phasing genotypes with ShapeIt5.1.0 (Loh et al. 2018; Loh et al. 2020; Hofmeister et al. 2023).

In addition to short-read sequencing, the TMM BirThree Cohort Project also conducted long-read sequencing to analyze structural variations (Otsuki et al. 2022). To ensure accurate assessment of deletions and validate SNV calls, we utilized long-read sequencing data from 109 trios that had already been analyzed using short-read sequencing. Briefly, genomic DNA from activated T cells was sequenced using nanopore technology via the ONT PromethION system. For SV detection, we followed the ONT protocol, employing LRA for alignment and CuteSV for SV calling (Jiang et al. 2020; Ren and Chaisson 2021).

### Quality control (QC) for extracting interparental gene conversion sites

Interparental gene conversion sites were identified based on Mendelian errors detected in the SNV calling data derived from short-read sequencing. This method requires stringent quality control because base calling or sequencing errors would otherwise increase the number of false positives. We developed QC filters based on previous studies that investigated de novo point mutations by focusing on Mendelian errors (Sasani et al. 2019; Bergeron et al. 2021). Target variant sites were restricted to biallelic SNVs on autosomal chromosomes.

### Sample-level filters

We calculated PI_HAT values between individuals using PLINK v2.0 and excluded trios for which the parent-child PI_HAT value was below 0.4. Individual the heterozygosity was assessed, and samples with the heterozygosity exceeding 34.5% were excluded due to concerns regarding cross-contamination.

### Site-level filters

Due to the difficulty in calling variants in low-complexity regions (LCRs), these areas were excluded from our analysis (Sasani et al. 2019). Variant sites with a call rate of 95% or lower were also excluded. We calculated the Hardy-Weinberg equilibrium P-value using founder individuals and excluded any sites with a P-value less than 10^-6.

### Variant-Level Filters

For each sample, we calculated the average depth and excluded variant sites in which the DP was below half or above twice the average. Variant sites with GQ scores < 60 for each individual were removed. Sites were excluded if two or more reads supported the alternative allele in the reference homozygote or the reference allele in the alternative homozygote. Given that somatic mutations, mapping errors, and sample contamination tend to peak at an allelic balance (AB) of approximately 20% (Besenbacher et al. 2015), sites with an AB below 30% or above 70% were removed.

### Data access

The WGS data generated in this study has been submitted to the NBDC Human Database under accession number hum0184.v2 and is available, at least in part, through controlled access (https://humandbs.biosciencedbc.jp/en/hum0184-v2).

## Acknowledgments

Author contributions: Jum.T. and Jun.T. performed the computational analyses. The Tohoku Medical Megabank Project Study Group contributed to the data collection. Jum.T. and Jun.T. wrote the manuscript. A.S., T.S., Y.T., H.H., E.Y., Z.W., M.S. and N.Y. designed the experiments. M.T., J.S., G.T. and Jun.T. conceived and supervised the study. The manuscript was reviewed, and valuable comments were provided by Shoukhrat Mitalipov and Aleksei Mikhalchenko, both from the Center for Embryonic Cell and Gene Therapy, Oregon Health & Science University. All the authors have read and approved the final version of the manuscript. This work was supported by the JSPS Grant-in-Aid for Scientific Research (KAKENHI) (C), Grant Number 21K09487.

